# The feature landscape of visual cortex

**DOI:** 10.1101/2023.11.03.565500

**Authors:** Rudi Tong, Ronan da Silva, Dongyan Lin, Arna Ghosh, James Wilsenach, Erica Cianfarano, Pouya Bashivan, Blake Richards, Stuart Trenholm

## Abstract

Understanding computations in the visual system requires a characterization of the distinct feature preferences of neurons in different visual cortical areas. However, we know little about how feature preferences of neurons within a given area relate to that area’s role within the global organization of visual cortex. To address this, we recorded from thousands of neurons across six visual cortical areas in mouse and leveraged generative AI methods combined with closed-loop neuronal recordings to identify each neuron’s visual feature preference. First, we discovered that the mouse’s visual system is globally organized to encode features in a manner invariant to the types of image transformations induced by self-motion. Second, we found differences in the visual feature preferences of each area and that these differences generalized across animals. Finally, we observed that a given area’s collection of preferred stimuli (‘own-stimuli’) drive neurons from the same area more effectively through their dynamic range compared to preferred stimuli from other areas (‘other-stimuli’). As a result, feature preferences of neurons within an area are organized to maximally encode differences among own-stimuli while remaining insensitive to differences among other-stimuli. These results reveal how visual areas work together to efficiently encode information about the external world.

## Introduction

Our visual system makes sense of the world via its myriad neurons, each tasked with extracting specific visual features from the environment. Decades of work in primates have shown that after visual inputs arrive in primary visual cortex (V1), they proceed to numerous higher visual areas (HVAs)^1^. At an inter-area level, each HVA is thought to prefer a specific portion of the visual world (e.g. there is a preference for orientated edges in V1^2^; there is a preference for faces in fusiform face area^3^). At an intra-area level, individual neurons exhibit diverse tuning preferences that enable encoding of the portion of the visual world that a given HVA is concerned with (e.g. V1 possesses neurons with preferences for edges of different orientations^4^; face-selective areas possess neurons preferring different aspects of faces^5^). However, due in part to the large size of visual areas in primates and the resulting difficulty in broadly recording from neurons within and across different HVAs, a clear understanding of how across-area differences in feature preferences relate to within-area organization of feature preferences is lacking. This limits our ability to gain a holistic view of the visual system and to understand how its computational goals guide its functional organization.

The mouse visual system provides a tractable model for addressing these issues. Anatomical tracing^6–9^ and functional mapping^10–13^ of mouse cortex have revealed ∼10 distinct HVAs, most of which receive significant direct input from V1. Circuit tracing and light-evoked spike time analyses have found some evidence that mouse HVAs are organized in a hierarchal manner^6,7,9,14^, although these studies have also found that mouse HVAs appear to be more strongly interconnected than those in primates^6–9^. In one study^14^, a single hierarchy was described: V1→LM→RL→LP→AL→PM→AM (for full nomenclature of mouse HVAs, see Methods). In contrast, other studies attempting to parallel primate visual streams have sorted the mouse hierarchy into putative ventral (V1→LM→P→LI→POR) and dorsal (V1→RL→AL→A→PM→AM) processing streams^7,9^. Additional studies have examined differences in visual feature preferences between mouse HVAs, but to date it has been difficult to draw firm conclusions. For instance, by presenting full field drifting grating stimuli, several studies have found differences in spatial and temporal frequency tuning properties between HVAs^10,11,15–17^, though some of the differences noted appear to arise from the specific inclusion criteria used^18^ or whether experiments were performed under awake or anesthetized conditions^19^. Another study^20^, which used 2-photon calcium imaging to record from different genetically-defined cell types, across different cortical layers and HVAs, is notable for revealing broad tuning curves and largely overlapping preferences within and across genetically-defined cell types, across cortical layers, and across HVAs. Thus, what remains missing is a) an inter-area level understanding of which portions of the visual world each mouse HVA is concerned with; b) an intra-area level understanding of how specific portions of the visual world are specifically encoded by neurons within each HVA; c) a cross-scale understanding of how inter-area feature preferences arise from and relate to intra-area feature preferences.

To address this, we used *in vivo* 2-photon calcium imaging to record from thousands of neurons across mouse V1 and five HVAs. We leveraged advances in modelling neuronal responses using artificial neural networks (ANNs)^21–23^ to build predictive models of the neurons, which we used to generate preferred stimuli for individual neurons. This enabled us to outline how mouse HVAs are functionally organized, ask why such an organization arises, and elucidate what the roles for such an organization are in encoding visual stimuli.

## Results

### Modelling multiple mouse visual cortical areas with artificial neural networks

To gain an understanding of the visual features encoded by neurons in various visual cortical areas in the mouse, we modelled neuron responses to visual stimuli in different areas with artificial neural networks (ANNs). We used this approach to determine the preferred stimulus (i.e. a visual stimulus that will strongly activate the neuron) for each neuron recorded *in vivo*, and to run *in silico* experiments that would be otherwise infeasible experimentally. To generate these models, we recorded from neurons in six visual cortical areas in mouse, including primary visual cortex (V1) and five HVAs spanning different anatomical hierarchical levels of both putative dorsal and ventral visual streams (LM, LI, POR, AL, and RL; **Figure 1a**). Each area was identified using widefield calcium imaging, with area segmentation based on the phase of retinotopic maps^12,13,24^, using transgenic mice in which the calcium indicator jRGECO1a was expressed throughout cortex^25^ (**Figure 1a**). After identifying a given cortical area, we used 2-photon calcium imaging to record light-evoked responses from neurons in layer 2/3 during the presentation of 2,500 different static natural images.

**Figure 1:**
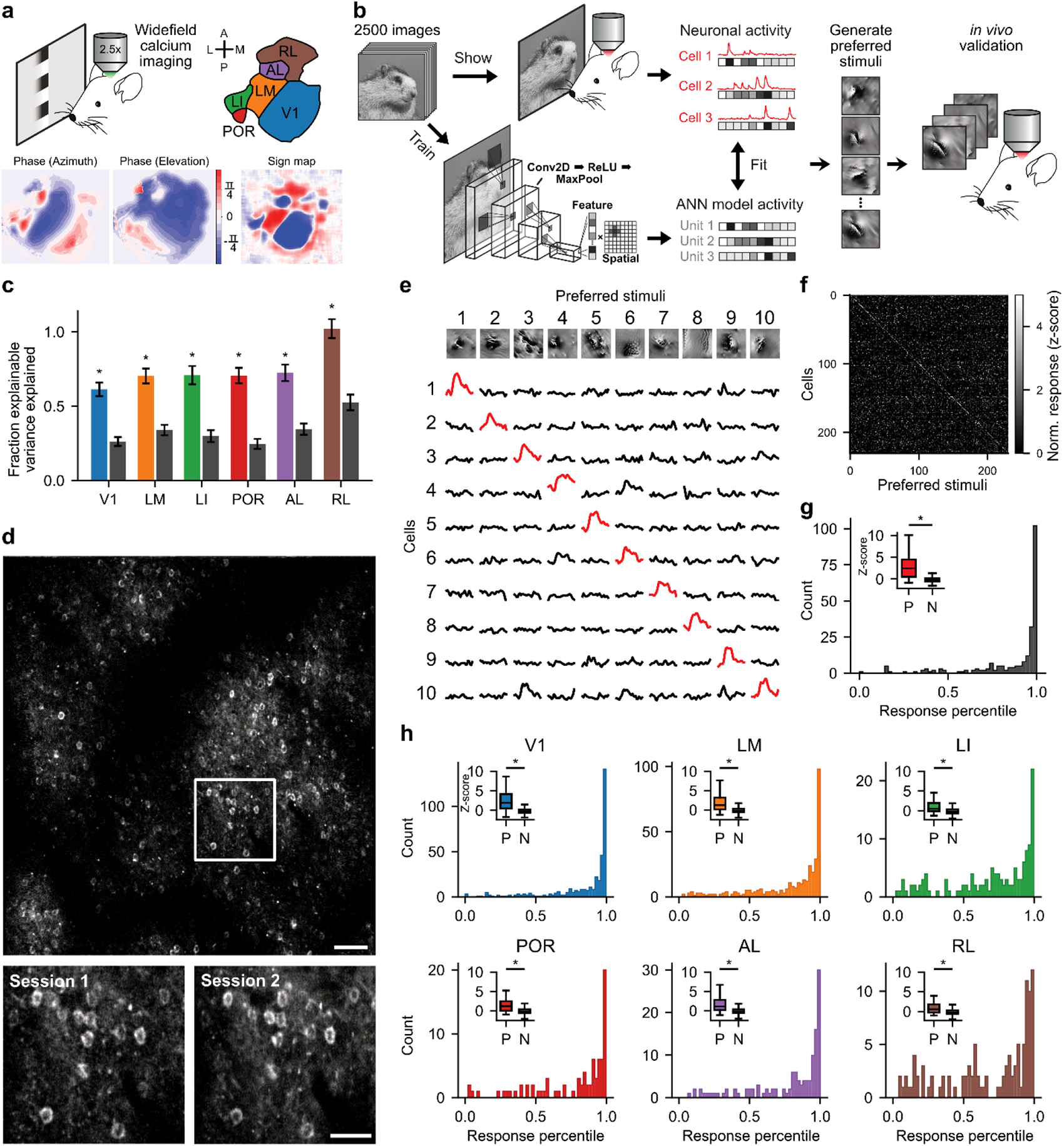
Modelling mouse visual cortex with artificial neural networks. **a**, Visual cortical areas were identified via retinotopic mapping using widefield calcium imaging. Six visual areas were targeted for subsequent 2-photon calcium imaging: V1, LM, LI, POR, AL, and RL. (A: Anterior, L: Lateral, M: Medial, P: Posterior). **b,** Workflow for building ANN models of mouse visual areas. Neuronal responses to 2,500 natural images were recorded in each area using 2-photon calcium imaging. For each area, an ANN model was trained to predict the neuronal responses to the same image set. We used the models to generate preferred stimuli. For validation experiments, preferred stimuli were presented back to the animal on a second day. **c,** Model performance measure as the fraction explainable variance explained for ANN models (*coloured*, left bars) compared to a linear model (*grey*, right bars). **d,** *Top*, Example 2-photon imaging field of view from a validation experiment. *Bottom*, Zoomed-in view of the region highlighted above. Scale bars: *Top*, 50 µm; *Bottom*, 30 µm. **e,** Responses of ten example neurons (rows) to their respective preferred stimuli (columns) from the experiment shown in (**d**). **f**, Same as (**e**), except for all 231 neurons simultaneously recorded in a single V1 session, showing a strong selective preference for preferred stimuli, with little ‘off-diagonal’ activity. **g,** For the recording session above, the distribution of response amplitudes of each neuron to its preferred stimulus, as a percentile of its response amplitude to all preferred and natural stimuli. *Inset* – Normalized neuronal response amplitudes to preferred (P) and natural (N) images. **h,** Same as (**g**), except pooling all validation experiments across all six visual cortical areas (V1: n = 364 neurons/2 mice, LM: n = 351 neurons/2 mice, LI: n = 129 neurons/3 mice, POR: n = 80 neurons/2 mice, AL: n = 126 neurons/2 mice, RL: n = 105 neurons/2 mice). All asterisks, p-value < 1e-11.

Next, we used ANNs to model the responses from neurons in each cortical area (**Figure 1b**). We used a shallow convolutional neural network with a factorized readout layer^26^ that separates the visual features (i.e. ‘what’) and the spatial locations in the images that drive neurons (i.e. ‘where’), referred to as the ‘spatial mask’. For each cortical area, we trained a unique ANN to predict the responses of individual neurons to the natural images. For model training, we selected neurons that reliably responded to natural image presentation (**Supplementary Figure 1**; see Methods). Additionally, only model units (i.e. digital twins of our real neurons) with > 30% explainable variance explained were used for subsequent analyses. This resulted in > 7,250 model units included for subsequent analyses (V1 = 1,418 units; LM = 1,469 units; LI = 1,112 units; POR = 1,070 units; AL = 1,298 units; RL = 899 units). Importantly, each ANN model had a significantly higher fraction of explainable variance explained compared to a simple linear model (**Figure 1c**; see Methods).

Finally, to validate our modelling strategy, for a subset of recordings (n = 2-3 animals per cortical region), we re-recorded from the same neurons on a second day while presenting the animal with a subset (n = 300-500) of the natural images shown the first day, as well as their preferred stimuli (**Figure 1d**). Each preferred stimulus, generated to maximally activate the twinned model unit, also strongly and selectively activated its corresponding *in vivo* neuron, with much smaller ‘off-diagonal’ activation compared to ‘on-diagonal’ activation (**Figure 1e-g**). The median response generated for all preferred stimuli was greater than 90% of the responses evoked by natural images (**Figure 1h**). Our results indicate that these ANNs can serve as tools for predicting neuronal activity and exploring visual feature preferences in areas V1, LM, LI, POR, AL, and RL.

### The functional organization of mouse visual cortex

We first examined the overall organization of visual cortex. To do so, we examined the functional similarity between HVAs by using our ANN models to test how features of visual stimuli were differentially represented across brain areas. We took advantage of the fact that we could align the spatial masks (the ‘Where-layer’) of the factorized readout layer for each model unit (**Figure 2a**), which allowed us to focus exclusively on differences arising from the ‘What-layer’. We presented 10,000 natural images taken from ImageNet^27^ to the spatial mask centered units. For each image, we measured the evoked population response (i.e. how strongly each model unit was activated by each image) and repeated this process for ANN models of V1 and the five HVAs. We then compared the similarity of population responses to the same set of natural images across visual areas by calculating the distance correlation (dCor), a non-linear method for measuring statistical dependence between two multivariate random variables (**Figure 2a**). We found that the distance correlations between pairs of HVAs showed clear differences in their strength (**Figure 2b**). However, we also observed that distance correlation values were relatively high between all areas, suggesting the presence of some widely shared correlations. To specifically focus on unique correlations shared between each pair of regions, we calculated the partial distance correlation (pdCor) between each pair of regions after conditioning out correlations with all other regions (**Figure 2a**). Focusing on pdCor, we found that activity in V1 was most similar to LM and LI. Activity in LM was most similar to RL and AL. LI and POR were most similar to each other, RL and AL were most similar to each other, and these latter two groups (LI+POR vs. RL+AL) were particularly dissimilar from one another (**Figure 2c**). We used multidimensional scaling (MDS) to visualize the distance correlations and partial distance correlations between the different areas as undirected graphs. This visualization emphasized the existence of two distinct functional streams emerging from V1 and passing through LM: one being V1→LM→LI→POR, the other being V1→LM→RL→AL (**Figure 2b,c**; note that since we know *a priori* that V1 is the predominant input region receiving signals from the visual thalamus, we orientated the MDS graph to place V1 at the bottom). Remarkably, a proposed dorsal/ventral two-stream hierarchy based on anatomical connectivity^9^ is in strong agreement with our results (**Figure 2d**), revealing a tight relationship between function and anatomy.

**Figure 2:**
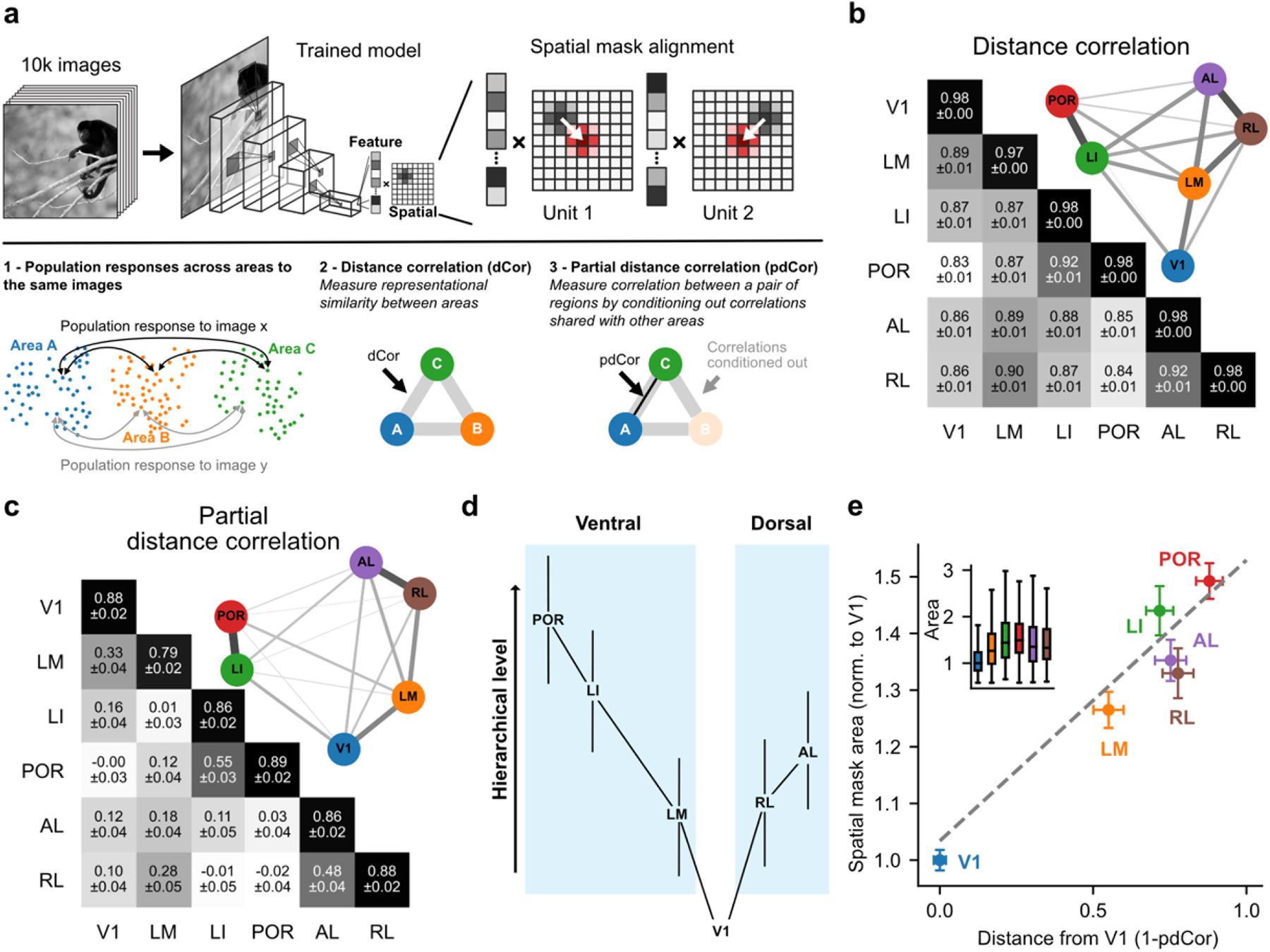
The functional organization of mouse visual cortex. **a,** *Top*, The spatial mask layer of each model unit was aligned, and each ANN was presented with the same 10,000 natural images. *Bottom*, Distance correlation (dCor) was used to compute the functional similarity between responses of pairs of areas. Partial distance correlation (pdCor) was used to condition out correlations shared globally across all areas. **b,** A matrix showing the pairwise distance correlation (dCor) between the stimulus manifolds of pairs of areas. *Inset*, Visualization of the resulting network structure using multidimensional scaling (MDS). **c,** Same as (**b**) but for pdCor. **d,** Two-stream hierarchy proposed by D’Souza et al. based on anatomical connectivity^9^. **e,** The functional distance from V1, defined as 1 – pdCor, is shown against the median spatial mask area of each visual area. *Inset* – Distribution of spatial mask area for each ANN model. For full statistical comparisons, see **Supplementary Figure 2**.

Finally, previous work has shown that receptive field size increases as one moves up the visual hierarchy^7,9,14,20^. We examined whether the spatial mask size in the factorized readout layer, which is loosely analogous to a receptive field, showed differences across visual cortical areas. We normalized mask size to V1 and found that spatial mask size increased along the visual hierarchy (**Figure 2e**). Overall, these analyses provide strong evidence that, on a functional level, mouse HVAs are representationally organized along two distinct hierarchical processing streams. Moreover, representations of natural images increasingly differ between HVAs as a function of hierarchical distance.

### Visual feature similarity with invariance to self-motion related image transformations guides the organization of feature preferences across HVAs

We next asked why HVAs are organized in this manner. Unlike in primates – where it is well-established that IT cortex is dedicated to processing visual objects^28^, and thus specific hypotheses can be formulated around how preferences for specific visual categories are organized in different HVAs^29–31^ – we did not have a clear *a priori* assumption for the type of model that might guide the organization of visual feature preferences in mouse cortex. We therefore decided to compare three simple image-based organizing models: a) a template-matching based model; b) a spatial frequency based model; c) a model based on visual feature similarity with invariance to image transformations that could arise as an animal moves around the world (we focused on invariance to affine transformations, including translation, scaling and in-plane rotation, as these are simple to apply to 2-D images).

To compare between these three models, we pooled the preferred stimuli from all areas. Next, for each model we constructed a 256-dimensional embedding space. For the template-matching based model, we performed a principal component analysis (PCA) on the preferred stimuli and defined the resulting principal component space as the embedding. For the spatial frequency based model, we performed a fast Fourier transform (FFT) on each preferred stimulus, followed by PCA. For the third model, we embedded the preferred stimuli using SimCLR, a contrastive deep learning tool specifically designed to group together images that are similar to one another^32^, and which has previously been used to measure the similarity of AI-generated preferred stimuli from primate V4^33^. SimCLR maximizes invariances towards arbitrary transformations of input images, which we achieved by applying a set of affine transformations to the preferred stimuli – translation (± 10% X/Y shift), rotation (± 90°), and resizing – then projecting these images into SimCLR’s latent space, and minimizing the distance between different transformations of the same preferred stimuli (**Figure 3a**). Notably, the transformations we applied are similar to those that would be expected from self-motion.

**Figure 3:**
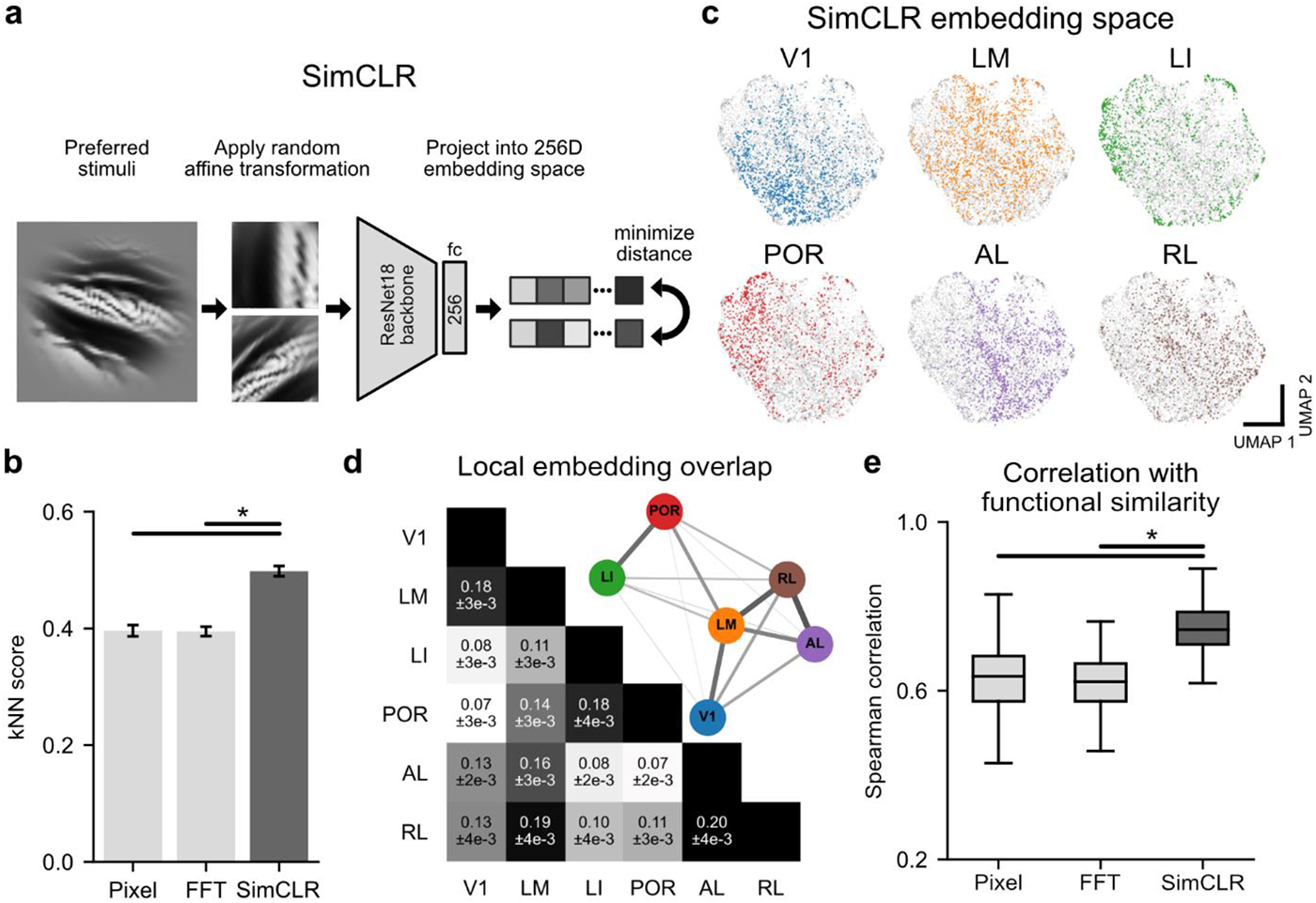
A preferred stimulus embedding based on image similarities with invariance to self-motion related transformations best explains the functional organization of mouse visual cortex. **a**, SimCLR was used to generate an embedding space that is invariant to affine transformations. The SimCLR model was trained to minimize the distance in its 256-dimensional embedding space between pairs of randomly transformed versions of a given preferred stimulus. **b,** Classification accuracy of stimulus labels using a kNN (k=20) trained either on a template-matching embedding (Pixel), spatial frequency embedding (FFT), or SimCLR embedding. **c,** Visualization of SimCLR’s embedding of all preferred stimuli, reduced to two dimensions using UMAP. Preferred stimuli from each area are colour-coded according to the visual area that each stimulus originated from. **d,** A matrix indicating the extent of local overlap between the stimulus manifolds in the SimCLR embedding space between each area, and the resulting network structure visualized using MDS. **e,** Spearman correlation of the pairwise overlap matrix (**d**) with the partial distance correlation matrix (Figure 2c) for Pixel, FFT, and SimCLR embeddings.

First, we tested which of these models was best able to group together preferred stimuli according to the area that they were generated from. For each embedding model of preferred stimuli, we performed a k-nearest neighbours analysis (kNN; k = 20) and examined its classification accuracy (**Figure 3b**). We found that the SimCLR method significantly outperformed both template-matching (pixel-level PCA) and spatial frequency (FFT) based models (**Figure 3b**), meaning that SimCLR was better at grouping together preferred stimuli from the same HVA. We thus further explored details of the SimCLR embedding of preferred stimuli.

We visualized SimCLR’s 256-dimensional space by projecting it down to two dimensions using UMAP^34^. In this visualization (**Figure 3c**), each point represents a preferred stimulus, and the distance between any two points indicates the relative similarity between those two preferred stimuli in SimCLR’s latent space: specifically, distance in this space relates to image similarity with invariance to affine transformations. While the UMAP plot did not reveal distinct clusters – as predicted by the strong pairwise functional correlation between areas (**Figure 2b**) – colour-coding the embedding according to the visual area that each preferred stimulus came from revealed strong biases in the extent to which preferred stimulus space was represented by each visual area (**Figure 3c**).

How do we know if SimCLR’s embedding of preferred stimuli is related to how the brain organizes visual feature preferences? To ask this question, we examined whether SimCLR’s embedding of preferred stimuli could recapitulate the functional organization of HVAs (**Figure 2b,c**). We reasoned that the functional relationship between HVAs could be reflected in the overlap of their respective manifolds within the SimCLR embedding space. To calculate this, for each preferred stimulus from each HVA, we measured the likelihood of finding preferred stimuli from other HVAs within its 20 nearest neighbours (**Figure 3d**). As visualized with an MDS plot (**Figure 3d**), SimCLR’s embedding of preferred stimuli resulted in an organization of HVAs that was remarkably similar to that generated from neuron population responses (**Figure 2b,c**). To quantify this, we measured the topological similarity between SimCLR’s embedding (**Figure 3d**) and the results from the partial distance correlation analysis (**Figure 2c**) via the Spearman correlation coefficient of the respective similarity matrices. This indicated a significantly higher topological similarity between the SimCLR embedding and the functional organization of HVAs compared to when the same analysis was run on the template-matching or spatial frequency based models (**Figure 3e**). These results suggest that, at a cortex-wide scale, HVAs encode visual features agnostic to their orientation, scale, and position, in line with a general invariance to types of image transformations that can be induced by self-motion.

### Each HVA prefers distinct visual features

What do the preferred stimuli for different HVAs look like, and how do they differ between areas? To address these questions, we visualized the two-dimensional UMAP projection of the SimCLR embedding. We split the embedding into a 40 x 40 grid, and for each tile in the grid we randomly visualized an image contained within that tile (**Figure 4**). We refer to this visualization as the ‘Feature landscape of visual cortex’. It can be seen from this visualization that SimCLR effectively sorted preferred stimuli based on image similarity, as neighbouring preferred stimuli appear alike. To examine visual feature preference differences across HVAs, we selected the 200 most representative preferred stimuli for each visual area, on which we performed additional analyses. This was done within the SimCLR embedding by using a kNN analysis (k = 100) to select the images with the most preferred stimuli from the same visual area within their neighbourhood.

**Figure 4:**
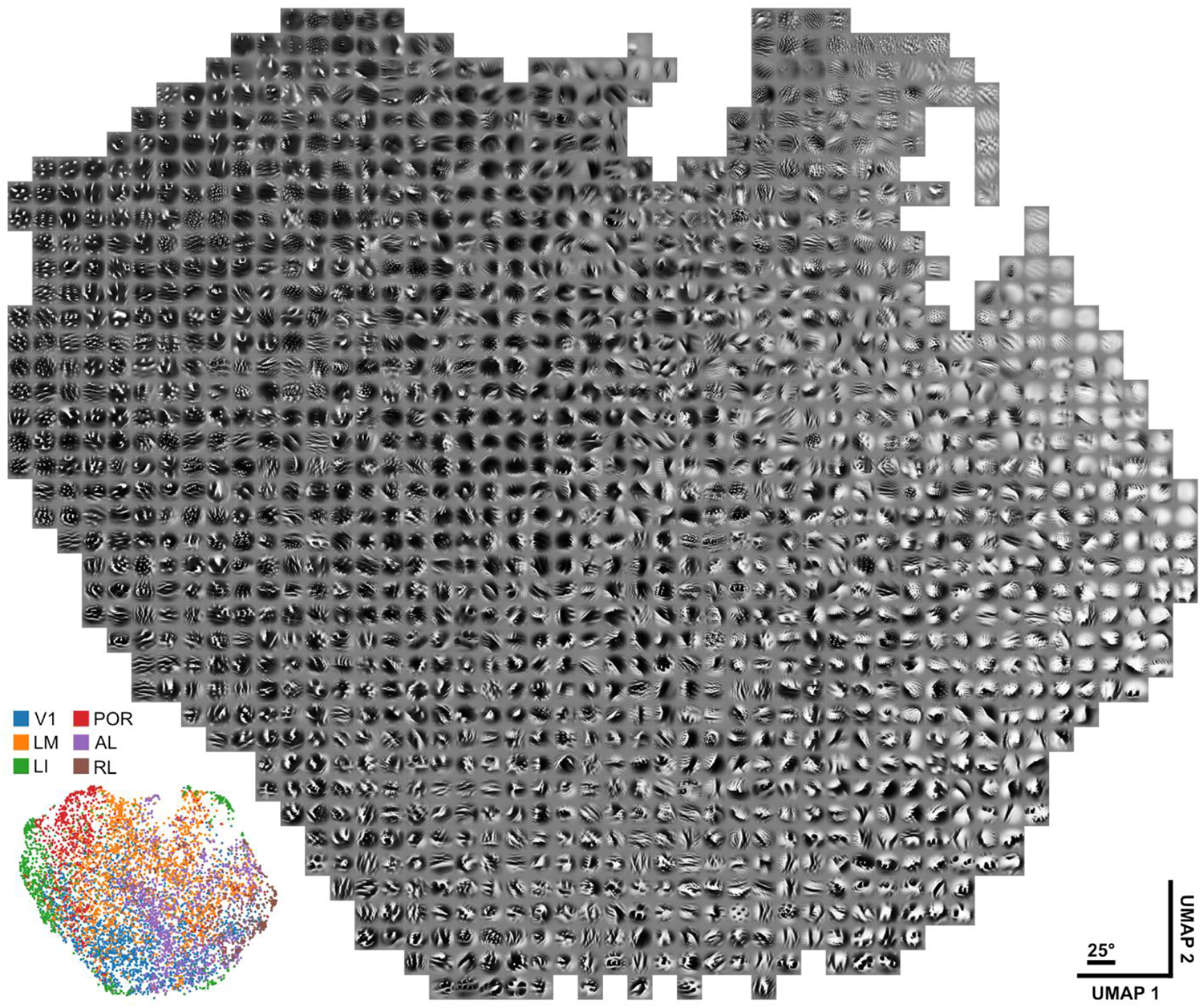
The feature landscape of visual cortex. Preferred stimuli were first projected into SimCLR’s 256-dimensional embedding space, then further projected onto a two-dimensional plane using UMAP. The UMAP projection was tiled into a 40 x 40 grid, and for each tile a random preferred stimulus contained within that tile is shown. *Inset* – Distribution of preferred stimuli from different cortical areas across the UMAP embedding (related to Figure 3c).

We observed several differences between the representative stimuli from different HVAs (**Figure 5a**). First, we noted differences in mean luminance, with some areas preferring darker or brighter stimuli. We found that preferred stimuli from LI and POR were particularly dark, whereas RL preferred stimuli were relatively brighter (**Figure 5b; Supplementary Figure 3**). Second, we noted differences in how many distinct segments made up preferred stimuli from each area. Calculating individual segments (measured via thresholding preferred stimuli into black and white segments), we found that preferred stimuli from most areas were best described as having white segments on top of a black background (i.e. significantly more white than black segments (except for AL)), and having smaller white segments than black segments (**Figure 5c,d**; **Supplementary Figure 3;** with LI having significantly more white segments than the other areas). Third, we noted that representative stimuli from some areas were dominated by lower spatial frequencies, whereas others possessed higher spatial frequency content. Performing an FFT and averaging radially over all orientations, we found that POR was dominated by the lowest spatial frequencies (though with increased power again at very high spatial frequencies), LM, AL, and RL were dominated by medium-to-low spatial frequencies, V1 was dominated by medium-to-high spatial frequencies, and LI was dominated by high spatial frequencies (**Figure 5e**; **Supplementary Figure 3**). Lastly, we noted that whereas areas V1, AL, and RL had representative stimuli that tended to feature long edges orientated along a single axis, areas LI and POR contained dotted segments that appeared to be arranged in grid-like patterns. Calculating folio symmetry (i.e. folding the image once) vs. quarto symmetry (i.e. folding the image twice) revealed that areas V1, LM, AL, and RL exhibited higher folio symmetry (**Figure 5f**; **Supplementary Figure 3**), whereas areas LI and POR exhibited significantly higher quarto symmetry (**Figure 5g**; **Supplementary Figure 3**). Importantly, these differences in image statistics for preferred stimuli from different HVAs persisted when we performed the same analyses on all preferred stimuli from each HVA, not just on the 200 most representative stimuli (**Supplementary Figure 4**). These results show that each HVA exhibits distinct preferences in visual features.

**Figure 5:**
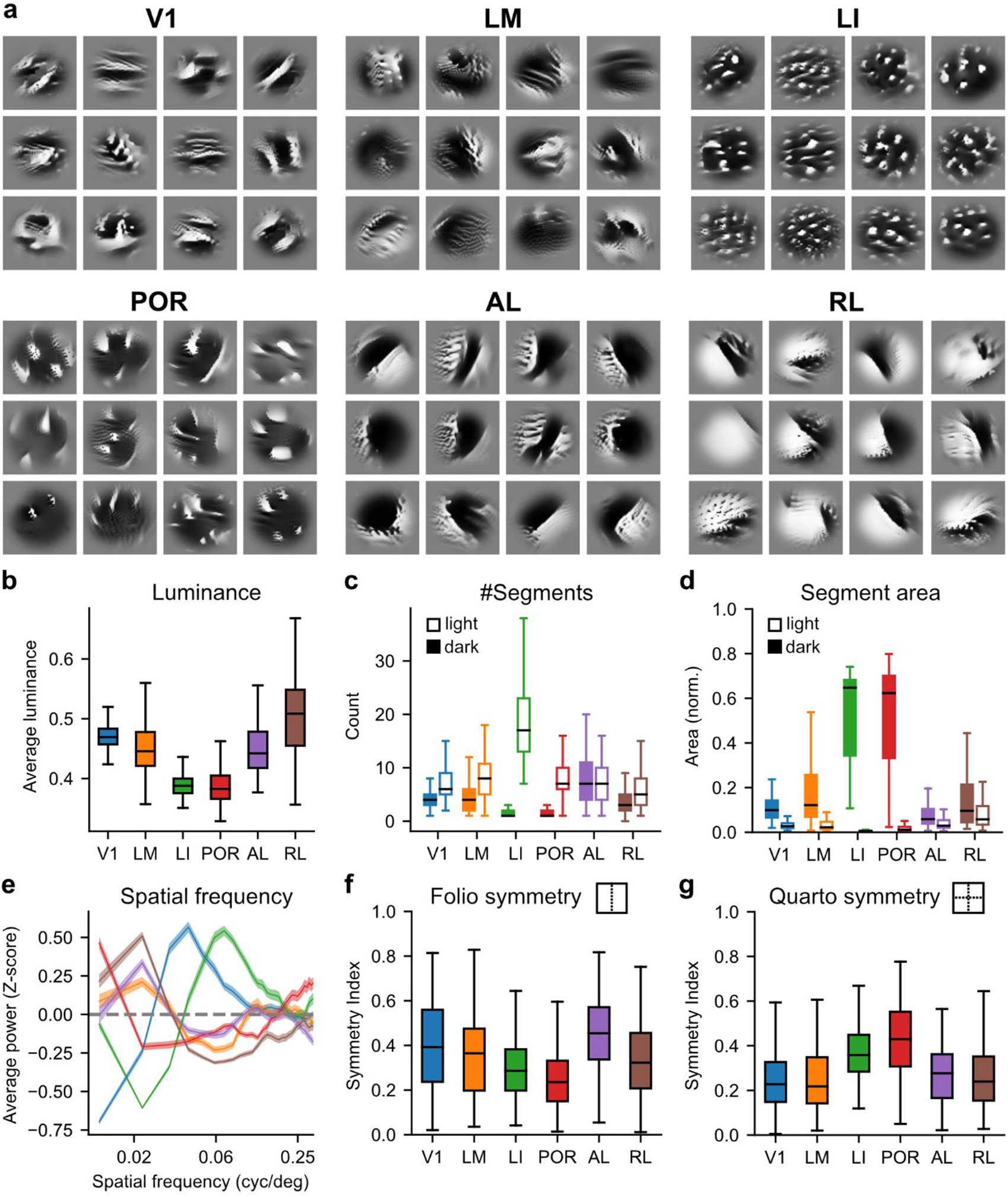
Mouse visual areas have distinct visual feature preferences. **a,** The most representative stimuli for each visual area. **b,** Mean luminance of representative stimuli across regions. **c,** Number of light and dark segments. **d,** Area of light and dark segments normalized to the area of the full stimulus. **e,** Radially averaged spatial frequency power spectrum. **f,** *Folio* (one-fold), and **g,** *quarto* (two-fold) symmetry index. For the full set of statistical comparisons, see **Supplementary Figure 3**.

### The set of preferred stimuli in a visual area represents a spanning set that effectively drive neurons through their dynamic range

The results above indicate that the preferred stimuli for each visual area possess distinct image statistics. As such, we wondered whether at a population level the set of preferred stimuli generated from a given area (which we refer to as ‘own-stimuli’) would drive stronger activity in the neurons from the same area compared to the set of preferred stimuli generated from other visual areas (‘other-stimuli’). To test this, we performed *in vivo* widefield calcium imaging with a new cohort of mice that were not included in the models from which we generated the preferred stimuli (**Figure 6a**). We presented a mix of preferred stimuli from all areas and found that, for all areas other than LM, own-stimuli drove significantly stronger activity in the area that they were generated from compared to other areas (**Figure 6b**; for *in silico* results, see **Supplementary Figure 5**). As such, the biases we found in feature preferences for each area are consistent and generalize across animals.

**Figure 6:**
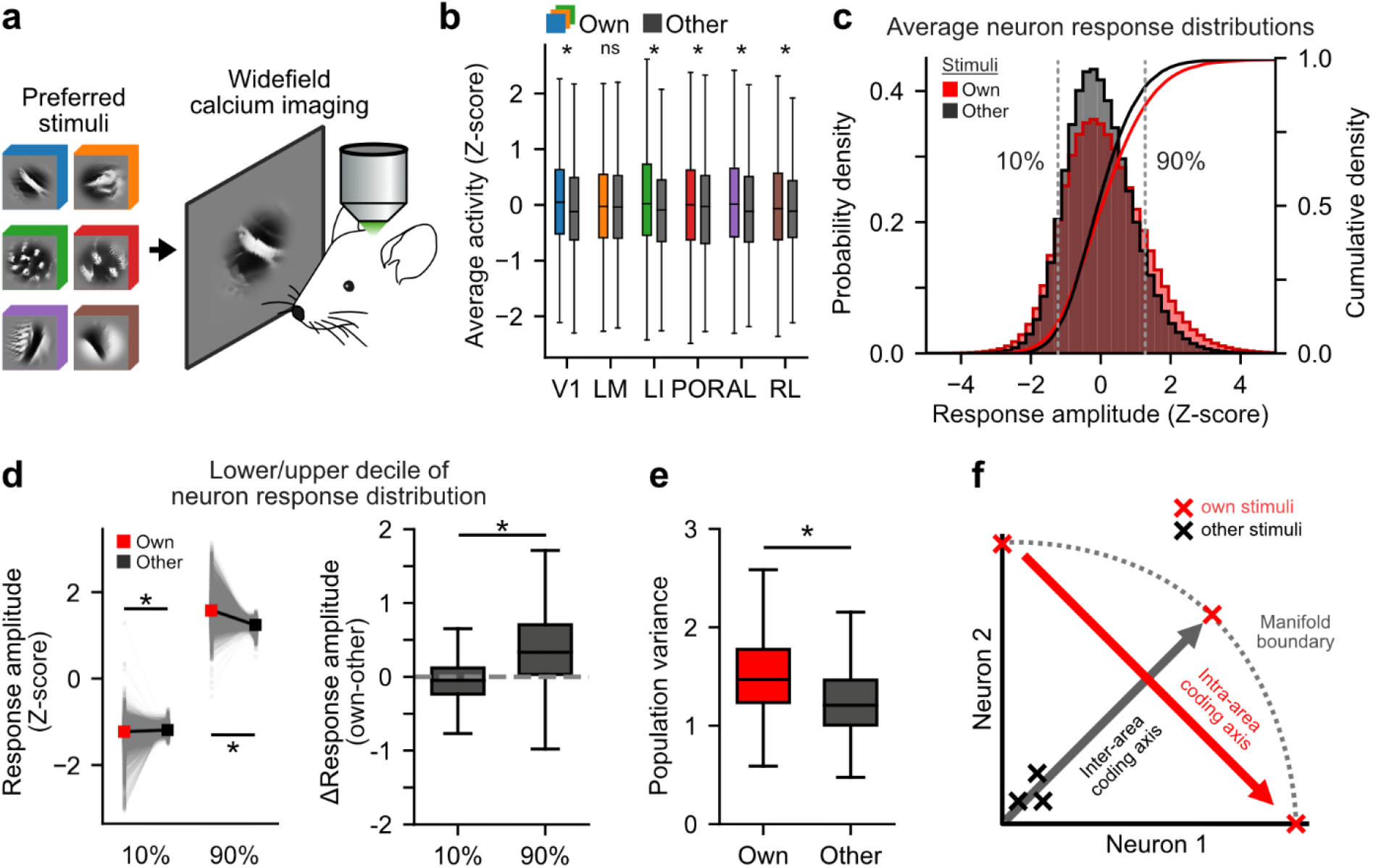
Intra-vs. inter-area feature coding axes arise from distinct trajectories through the neuron population manifold. **a**, Schematic outlining that preferred stimuli from various areas were presented to mice while activity across visual areas was measured using widefield calcium imaging. **b**, For experiments outlined in (a), average response amplitude within an area to its own preferred stimuli was compared to the activity that the same preferred stimuli drove in other areas. Wilcoxon rank sum test: for V1, LI, POR and AL, p < 0.001; for RL, p = 0.0118; for LM, p = 0.1007. See also **Supplementary Figure 5**. **C**, Example *in silico* data of response amplitudes of individual neurons to own-and other-stimuli, pooled across model units and all visual areas. *Left*, Probability density plots (histograms) of response amplitudes for all model units, pooled across all areas, evoked by the sets of ‘own-stimuli’ or ‘other-stimuli. *Right*, Cumulative density plots (line plots) of the same data. **d**, *Left*, The response amplitude (from *in silico* models, averaged across all areas) at the 10^th^ and 90^th^ percentiles evoked by own-stimuli and other-stimuli. Wilcoxon rank rum test: p < 0.001. *Right*, The *Left* data replotted to show that the 90^th^ percentile is significantly more different between own-and other-stimuli than the 10^th^ percentile. Mann-Whitney U test: p < 0.001. **e**, For *in silico* experiments, for all areas averaged together, the population variance in an area was greater when it was shown own-compared to other-stimuli. Mann-Whitney U Test: p < 0.001. **f**, Schematic of trajectories through the neuron population manifold. X’s represent responses to 3 different own-(*red*) and other-stimuli (*black*). Own-stimuli are located on the boundary of the population response manifold (the space of all possible population responses). Trajectories along the boundary of the manifold, i.e. rotations of the population vector, encode intra-area feature differences (intra-area coding axes, *red*), whereas trajectories from the center outwards, i.e. scaling of the population vector, correspond to axes encoding the overall feature differences across areas (inter-area coding axes, *black*).

Do own-stimuli drive stronger average activity simply because they contain more of the overall visual statistics preferred by neurons in a given area? To test this, we leveraged our *in silico* models, where we could center both own-and other-stimuli on individual models units by aligning their spatial masks, and we examined the range of responses evoked by sets of own-vs. other-stimuli. We found that instead of increasing the average response amplitude to all stimuli, own-stimuli drove neurons through a wider dynamic range (**Figure 6c**). To quantify this, we compared response amplitudes at the 10^th^ and 90^th^ percentiles and found that, compared to other-stimuli, own-stimuli extended the dynamic range over which neurons responded, on both upper and lower bounds (**Figure 6d**). However, the effect size at the 90^th^ percentile was significantly stronger than at the 10^th^ percentile (**Figure 6d**), which explains why own-stimuli drive stronger average activity in the area they were generated from compared to other-stimuli (**Figure 6b**). This also indicates that each own-stimulus, though generated to maximize the response of a specific neuron, actually also drives weak activity in many other neurons from the same area. Similarly, responses to own stimuli are often weaker than responses to other-stimuli. Consistent with this finding, we found that population variance (the extent to which a given stimulus drives diverse responses in a population of neurons, defined as the inter-quartile range of the distribution of responses to a given stimulus) was significantly higher for own-vs. other-stimuli (**Figure 6e**). Thus, the different features preferred by neurons within a given area can be viewed as a ‘spanning set’ that maximally drives neurons from that area through their dynamic range.

## Discussion

Here we described the feature landscape of visual cortex. By combining widefield calcium imaging, 2-photon calcium imaging, and modelling with ANNs, we generated digital twins of thousands of neurons in six different cortical areas in mouse, including primary visual cortex (V1) and five higher visual areas (LM, LI, POR, AL, and RL). Our results outline the functional organization of mouse visual cortex and provide a detailed understanding of how visual feature preferences are arranged across HVAs, why they are arranged in this manner, and what the outcomes of this organization are on encoding of the visual world.

How is the mouse visual system organized? Using a data driven approach, based on similarities of population activity to natural image presentations, assessed with distance correlation, we identified two hierarchically organized processing streams: V1→LM→LI→POR and V1→LM→RL→AL. These two processing streams are corroborated by previous studies that split mouse HVAs into dorsal and ventral streams based on anatomical tracing^7,9^ and certain functional response properties^35^, and are consistent with studies in mouse which found that receptive fields become larger at successive stages in the visual hierarchy^7,9,14,20^. Our finding that most areas’ preferred stimuli are biased towards lower luminance values build upon work from other mammalian species which have indicated that more neuronal resources are dedicated to processing OFF vs ON visual stimuli^36^. Furthermore, our findings of differences in spatial frequency preferences for preferred stimuli from different HVAs are consistent with some previous studies, which for instance found that RL and AL tended to prefer lower spatial frequencies, and V1 and LI tended to prefer relatively higher spatial frequencies^10,11,16,35^. Nonetheless, similar to previous studies of mouse V1^23,37^, using an ANN-based methodology to generate preferred stimuli allowed us to reveal how the various feature preferences (e.g. luminance, spatial frequency, receptive field size, etc.) interact to generate each neuron’s specific preferred stimulus, in various HVAs. For example, this revealed that many neurons in LI and POR appear to have preferences for dotted, grid-like patterns. However, to what purpose the mouse visual system developed these specific visual feature preferences across its HVAs remains an open question. To facilitate further exploration of the feature landscape of mouse visual cortex, we generated a graphical user interface that can be accessed [here](http://www.trenholmlab.com/featurelandscape.html).

Why is the mouse visual system organized in this manner? We took inspiration from primate models, which have tried to explain why the primate ventral visual stream is organized into functional patches preferring specific aspects of the visual world (e.g. faces, bodies, colours, etc.)^29–31^. Unlike in primates, where the hierarchy of HVAs is well matched by ANNs trained on object categorization, this does not appear to be the case for mouse cortex^38^, so we did not focus on testing models based on the computation of object categorization. However, much like primates, mice wander around the world and thus need to visually recognize things in a manner invariant to the types of transformations their visual system experiences^29^. We leveraged an ANN model that allowed us to sort preferred stimuli based on image similarity^32,33^ that was trained to be invariant to specific image transformations. We focused on the types of transformations that could be induced by self-motion and that are applicable to 2-D images: translation, scale, and in-plane rotation. We compared this to the most naïve model possible, a template-matching model that compared pixel-level similarity between preferred stimuli, and another model based solely on spatial frequency content, which was the focus of many early studies of mouse HVAs^19^. Our results indicate that, of the models tested, the model based on image similarity with invariance to self-motion related transformations best explained how visual feature preferences are organized in the mouse’s visual cortex. This suggests that at a cortex-wide level, while HVAs have arisen to extract distinct visual features from the world, they have done so in a way to minimize the effect of the types of transformations that could arise from self-motion. Nonetheless, it should be noted that whereas our SimCLR model only contained affine transformations, future work that studies exactly how a mouse’s body and eyes move in tandem could generate a more ecologically relevant set of image transformations, and we predict that such a model would outperform the one we present here.

What are the implications of this functional organization on how the visual world is encoded by the mouse? We found that the answer depends on the scale at which the question is asked. At the brain-wide level, the visual system is organized to encode features in a manner that is invariant to self-motion related image transformations (**Figure 3**). In turn, neurons in a given area possess an overall set of visual feature preferences that distinguish one area from another. The differences in area-wide feature preferences are delineated in the feature landscape of visual cortex (**Figure 4**). Though the preferred stimuli from each HVA do not clearly cluster, their image statistics are significantly different from one another (**Figure 5**). Finally, at the intra-area level, we find that individual neurons are driven through their dynamic range to a greater extent when presented with the set of own-stimuli compared to other-stimuli (**Figure 6**). This occurs because even though each preferred stimulus was designed to strongly activate a specific neuron, own-stimuli drive weak activity in many other neurons from the same area.

Since own-stimuli maximally activate their target neurons, responses to own-stimuli can be thought of as lying at the boundary of the neuron population manifold (the space of all possible patterns of activity for a population of neurons). Trajectories through the neuron population manifold that drive neurons from low to high activity can then be thought of as tuning curves. Now consider trajectories along the boundary of the manifold (Figure 6f, red arrow). Our results show that these trajectories effectively drive individual neurons through their dynamic range. Since movement along the boundary corresponds to rotations of the population vector, neuron population activity is also strongly decorrelated along these directions through the manifold. As such, trajectories along the boundary represent an efficient way to encode differences between the tuning preferences of neurons in the same area (which we refer to as ‘intra-area coding axes’). In contrast, consider trajectories that move from the center of the manifold outwards towards the boundary – these correspond to scaling of the population vector (Figure 6f, black arrow). These trajectories encode information that is shared across the neuron population, such as transitioning from the overall features preferred by one visual area to those of another (which we refer to as ‘inter-area coding axes’). Notably, the strong correlations between neurons that arise when moving from manifold center towards the boundary mean that such trajectories fail to encode differences in preferred features of neurons within an area. Therefore, since preferred stimuli lie at the boundary of the manifold, traversal through the set of preferred stimuli provides a principled and data-driven way of constructing tuning curves for a population of neurons. This is especially useful for brain areas whose neurons exhibit highly mixed tuning preferences or in which tuning properties are otherwise not easily parameterized. Finally, future work is needed to develop methods for identifying which specific trajectories through the set of preferred stimuli represent ecologically meaningful coding directions.

## Acknowledgements

We thank E. Macé for assistance setting up HVA identification using widefield calcium imaging. We thank M. Krause, K. Kuchibhotla and C. Pack for helpful feedback on the manuscript. We acknowledge our funding sources: Jeanne Timmins Costello Fellowship and VHRN Recruitment Award Scholarship to R.T.; HBHL Fellowship to RdS; Vanier Scholarship to A.G.; NSERC CGS-D to D.L; HBP Special Grant Agreement 3 and BBSRC (BB/Y003020/1) to J.W.; NSERC CGS-M and FRQNT Scholarship to E.C.; NSERC Discovery Grant (RGPIN-2020-05105), NSERC Discovery Accelerator Supplement (RGPAS-2020-00031), Arthur B. McDonald Fellowship (566355-2022), and CIFAR Canada AI Chair (Learning in Machine and Brains Fellowship) to B.A.R.; HFSP Career Development Award, Sloan Research Fellowship, ONR-Global Research Grant, Canada Research Chair, and CIHR Project Grant to S.T. This research was also enabled in part by support provided by Calcul Québec (https://www.calculquebec.ca/en/) and Compute Canada (www.computecanada.ca). The authors acknowledge the material support of NVIDIA in the form of computational resources.

## Author contributions

Experiments were designed by R.T., R.d.S., D.L., A.G., B.A.R., and S.T. *In vivo* data was collected by R.T., R.d.S., and E.C. Data analysis and statistics were performed by R.T., R.d.S., and J.W. ANN design and modelling was performed by R.T., D.L., A.G., and B.A.R. P.B. advised on issues related to ANN design and implementation. Figures were generated by R.T. The paper was written by R.T. and S.T.

## Data availability

All processed data included in the manuscript will be made available upon publication.

## Code availability

The code for ANN modelling and analysis can be accessed from https://github.com/Trenholm-Lab/MouseFeatureLandscape.

---

**Supplementary Figure 1:**
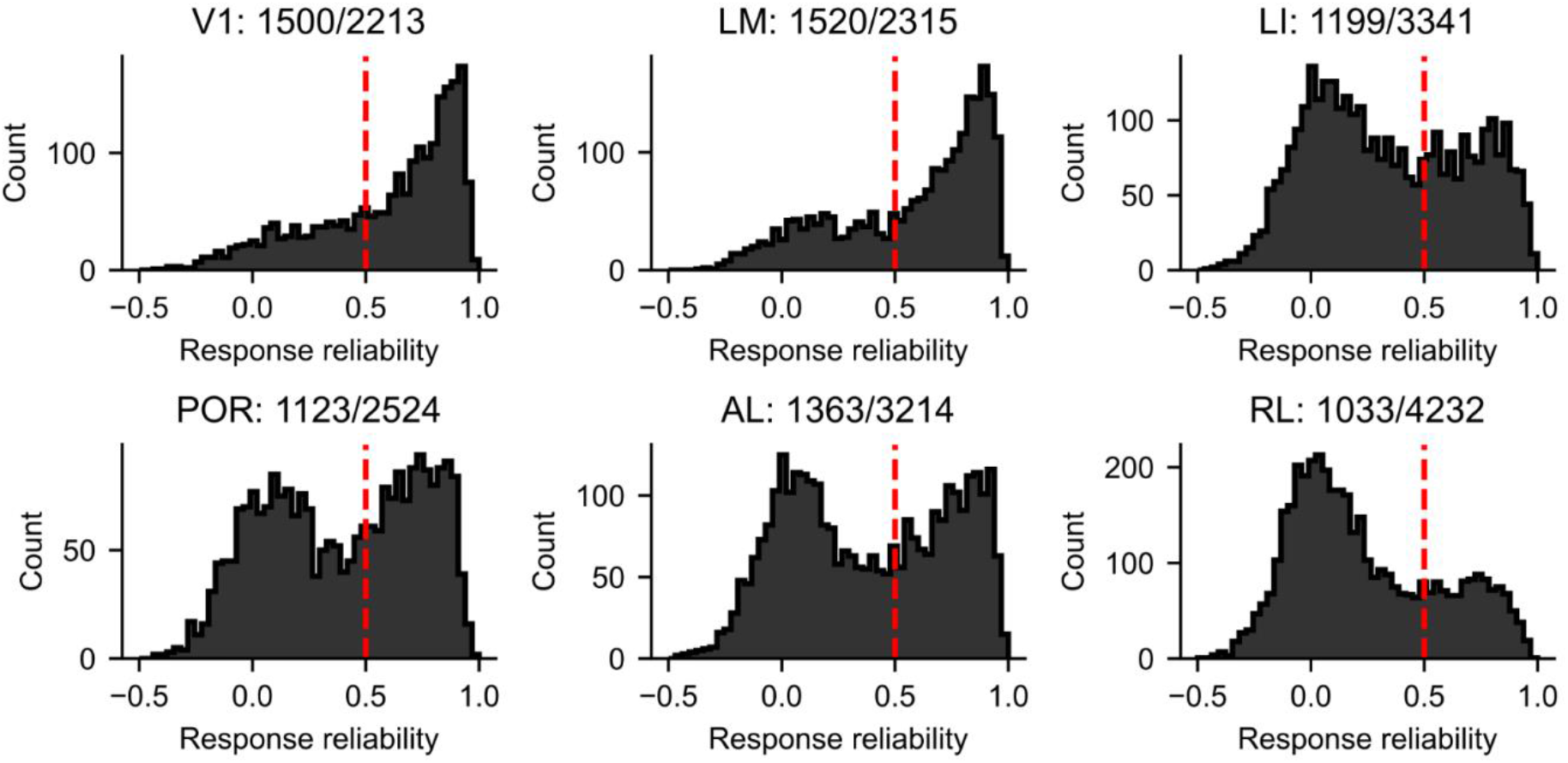
Distribution of response reliability of all neurons recorded across all areas. Response reliability was calculated as the Spearman-Brown corrected correlation coefficient across half-splits of repeated presentation of 100 natural images. Many areas exhibited a clear bimodal distribution. Neurons with reliability > 0.5 were used for subsequent ANN modelling (dashed red line). Numbers indicate the number of reliable neurons/total neurons.

**Supplementary Figure 2:**
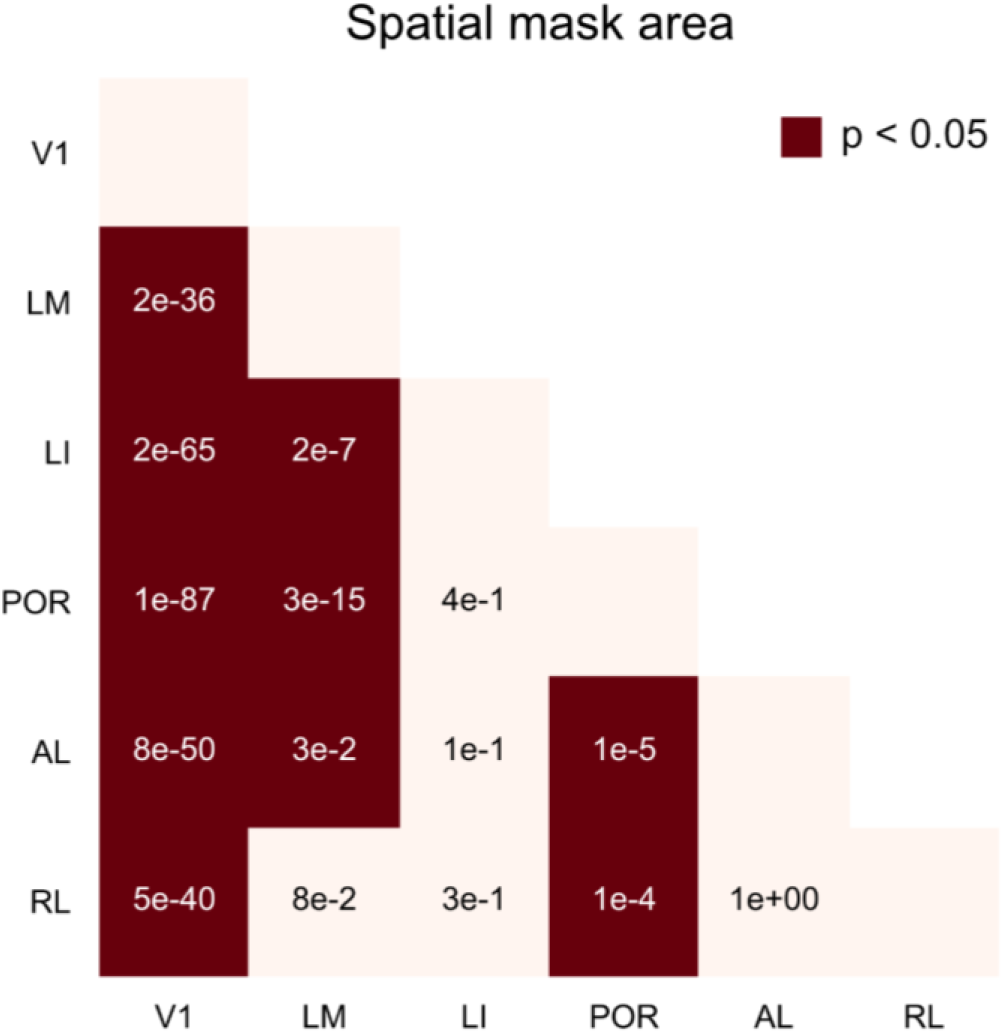
Pairwise statistical differences of spatial mask area (related to Figure 2e). Significance was assessed via Kruskal-Wallis and post hoc Dunn’s test with Bonferroni correction. Values indicate p-value, and significantly different pairs are highlighted in red. V1: n = 961 model units (67.7%), LM: n = 802 model units (54.6%), LI: n = 530 model units (47.7%), POR: n = 548 model units (51.2%), AL: n = 610 model units (47.0%), RL: n = 431 model units (47.9%).

**Supplementary Figure 3:**
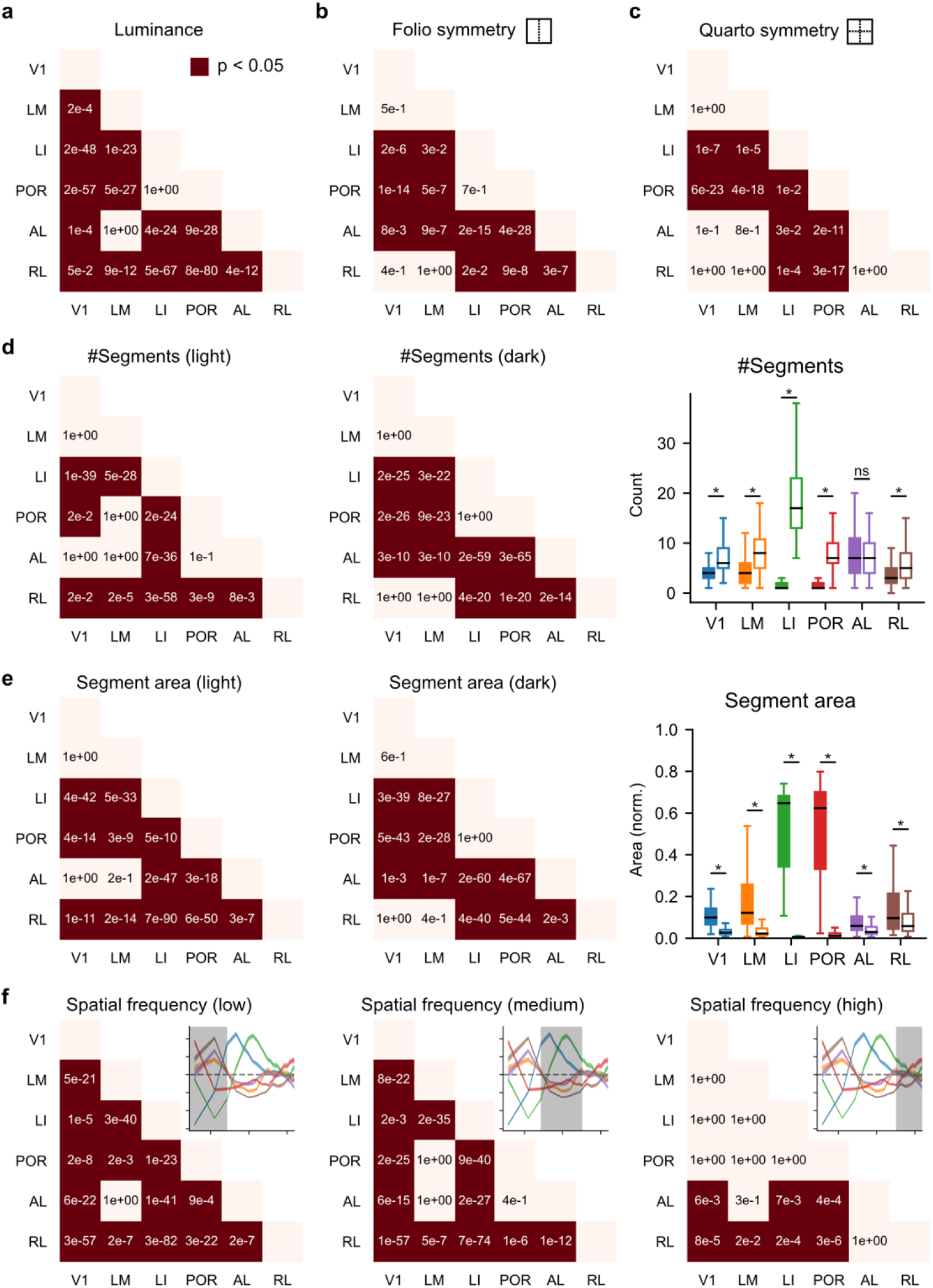
Pairwise statistical differences of visual feature preferences (related to Figure 5). Significance was assessed via Kruskal-Wallis and post hoc Dunn’s test with Bonferroni correction. Values indicate p-value, and significantly different pairs are highlighted in dark red. n = 200 stimuli/area **a**, Average pixel luminance. **b**, Folio symmetry index. **c**, Quarto symmetry index. **d**, Number of light (*left*) and dark (*middle*) segments, and difference between the light and dark segments within each area (*right*). **e**, Same as (d) but for segment area. **f**, Spatial frequency preferences were binned into low (0-0.04 cyc/deg), medium (0.04-0.15 cyc/deg), and high frequency bands (0.15-0.34 cyc/deg).

**Supplementary Figure 4:**
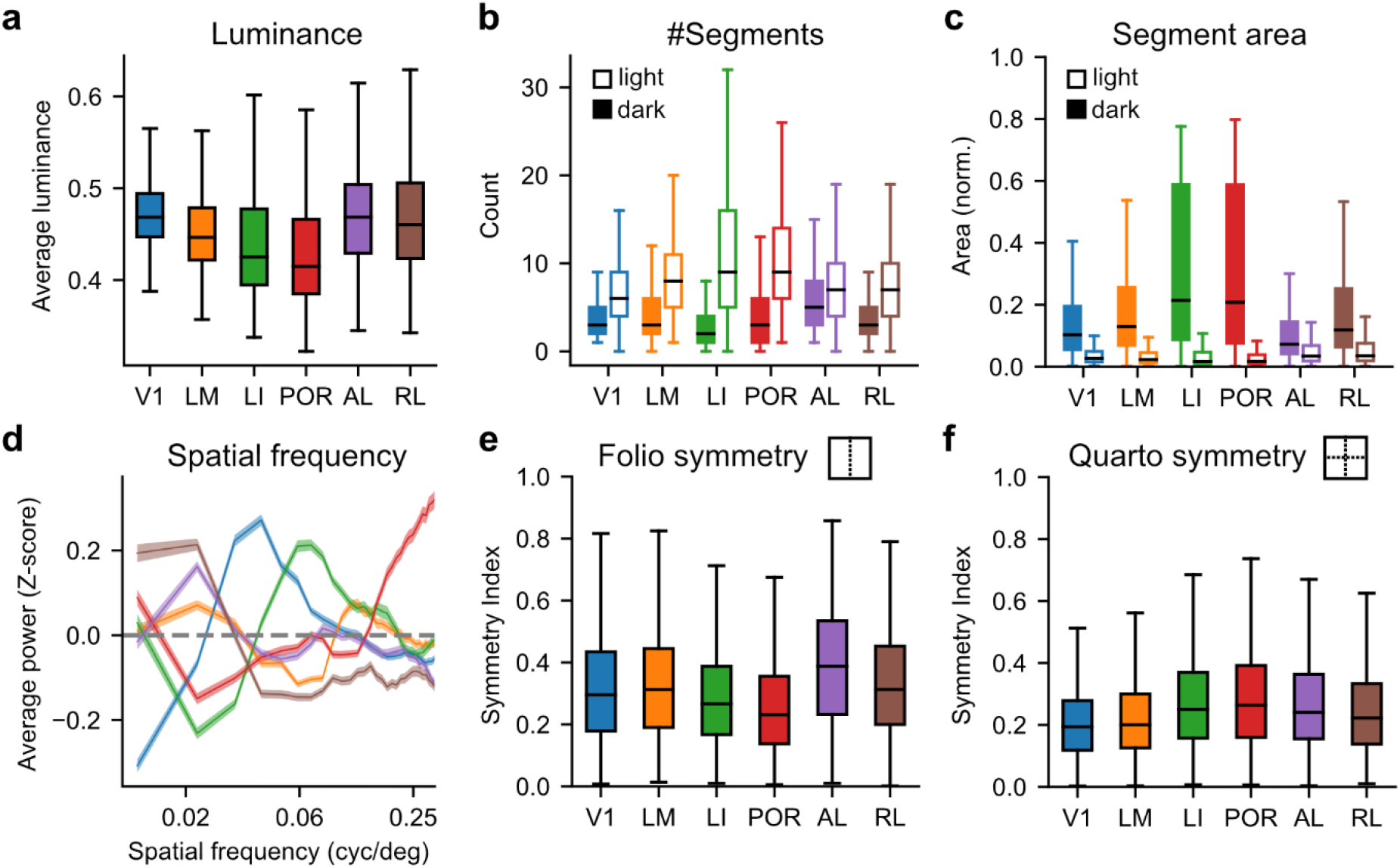
Visual feature preferences calculated across all preferred stimuli. The same as Figure 5b-g, except instead of calculating these metrics only on the 200 most representative stimuli for each HVA, here the metrics were computed for all preferred stimuli from each area. **a,** Mean luminance of preferred stimuli across regions. **b,** Number of light and dark segments. **c,** Area of light and dark segments normalized to the area of the full stimulus. **d,** Radially averaged spatial frequency power spectrum. **e,** *Folio* (one-fold), and **f,** *quarto* (two-fold) symmetry index.

**Supplementary Figure 5:**
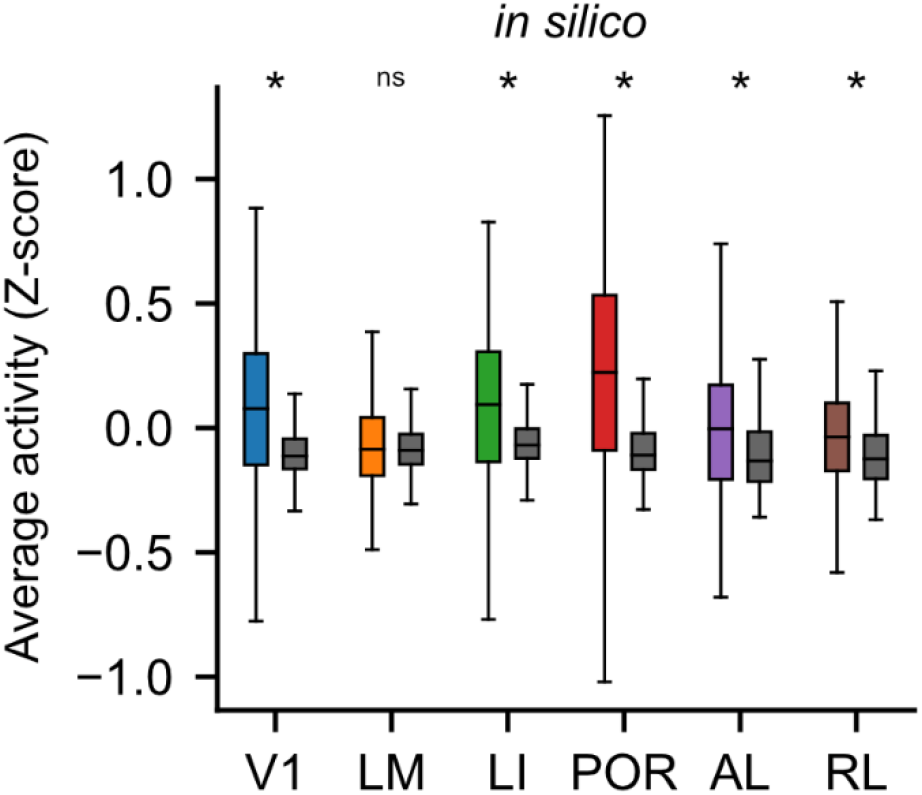
Own-stimuli drive increased area-wide activity in the area they were generated from. The same as Figure 6b, but from our *in silico* models. The much larger effect size for the *in silico* compared to *in vivo* results could arise from the fact that, unlike the *in silico* experiments, we are unable to align the stimuli to be centered over each cell’s receptive field *in vivo*, or it could arise because the effect size is small and swamped out by noise, or due to a combination of these issues. Wilcoxon rank rum test: for V1, LI, POR, AL and RL, p < 0.001; for LM, p = 0.0575.

**Supplementary Table 1:** Hyperparameters used for training ANN models.

| Area | Learning rate | Learning rate decay | L1 | L2 |
| --- | --- | --- | --- | --- |
| V1 | 0.001035 | 0.01986 | 0.0006565 | 0.0082100 |
| LM | 0.001562 | 0.04827 | 0.0002147 | 0.0006054 |
| LI | 0.001011 | 0.03148 | 0.0001245 | 0.0001245 |
| POR | 0.001404 | 0.04147 | 0.0019060 | 0.0019060 |
| AL | 0.001421 | 0.03263 | 0.0004504 | 0.0004504 |
| RL | 0.002319 | 0.02160 | 0.0052730 | 0.0052730 |

## Methods

### Animals

All procedures were performed in accordance with the Canadian Council on Animal Care and approved by the Montreal Neurological Institute’s Animal Care Committee. Mice used were Thy1-jRGECO1a-WPRE, line GP8.31 (The Jackson Laboratory #030526). All mice were adults (2-6 months old) and mice of both sexes were included. Mice were maintained in a temperature and humidity controlled facility on a 12 hr light/dark cycle.

### Head-bar and cranial window implantation

Mice were anesthetized with a cocktail containing fentanyl (0.05mg/kg), medetomidine (0.5mg/kg), and midazolam (5mg/kg)^39^. Skin was cut away over the skull, and a custom head-bar (adapted from a design from the Polley lab (Harvard University)) was attached to the skull over the right hemisphere using dental cement (C&B Metabond). Next, a 5 mm circular cranial window was made on the left hemisphere over visual cortex and sealed with a 5 mm glass coverslip (Warner) that was held in place with super glue. The exact position of the cranial window varied from mouse to mouse to enable easier optical access to the specific higher visual areas we were interested in.

### Widefield calcium imaging

Widefield imaging was performed similarly to previous studies^11–13,24^. In brief, an awake head-fixed mouse was placed under a 2-photon microscope (Neurolabware) with an independent epifluorescent imaging pathway. A 5X objective (Mitutoyo, M Plan Apo) was used to pass excitation light and collect emitted light. Excitation light was generated by a white LED (Thorlabs, MCWHL5), passed through a 559 nm excitation filter (Thorlabs, MF559-34), and a 588 nm dichroic (Thorlabs, MD588). Emission light passed back through the dichroic and a 630 nm emission filter (Thorlabs, MF630-39) and was captured at 10 Hz by a digital camera (PCO edge 3.1 M). We recorded light-evoked calcium responses through the cranial window on the left hemisphere, while visual stimuli were presented to the right eye. The mouse was presented with an inverting checkerboard stimulus that passed in both directions along both azimuth and elevation (10 repetitions for each direction) on a 24-inch computer monitor (BenQ RL2455) positioned 13 cm from the mouse’s eye. The moving stimulus was a 20° wide bar that was periodically swept across the monitor at a velocity of 4 °/s. The bar was filled with a checkerboard pattern (25° spatial frequency) reversing at 6 Hz. Spherical stimulus correction was applied to compensate for the flatness of the monitor^11^. The video data was first high pass filtered at half the stimulus frequency. A discrete Fourier transform (DFT) at the stimulus frequency was then applied:

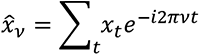

where ν is the stimulus frequency, and the phase difference between directions offset by 180 degrees was calculated to correct for the response delay due to slow dynamics of the calcium dye:

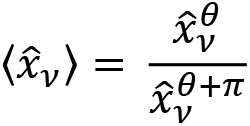

where *θ* is the direction of the stimulus. Phase and amplitude were then extracted. To generate a sign map, the difference between the gradients of the phases for both azimuth and elevation stimuli was calculated:

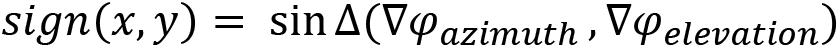

To identify HVA boundaries, the sign map was standardized, thresholded at 1.5 times the standard deviation, and denoised via binary opening. Lastly, a few (1-4) iterations of binary erosion were applied to refine the boundaries of each region.

For widefield experiments in Figure 6, static images were presented for 0.5 s with an inter-stimulus interval uniformly distributed between 1.3-1.7 s. Calcium activity was averaged for each visual area, identified by retinotopic mapping. The data was then high pass filtered at half the stimulus frequency, normalized to baseline, and denoised using singular value decomposition (SVD). Specifically, we found that the first right singular vector of the data matrix (stimulus x time matrix) corresponded well to the stimulus-evoked response kernel, and we therefore defined the response amplitude as the projection of the data matrix onto this vector. In order to compare whether a given stimulus more strongly activated neurons in the area it was generated from, we compared its response amplitude against the average amplitude in all other areas.

### 2-photon calcium imaging

2-photon imaging was performed similarly to previously described^40^. In brief, the laser (Insight X3, Spectra-Physics) was set to 1080 nm, and head-fixed, awake animals were placed under a resonant-galvo scanning 2P microscope (Neurolabware). Recordings were acquired at 10 Hz, from neurons in layer 2/3 (depth between 120 and 300 µm measured at the center of the FOV). Animals were presented with 2500 natural images from ImageNet^27^. Images were scaled to a size of 135 x 135 pixels, converted to grayscale, and shown for 0.5 s with a vertical height of 98 degrees. The inter-stimulus interval was uniformly distributed between 1.3-1.7 s so that responses were not entrained by a fixed stimulation frequency. A subset of the images (100/2500) were repeated 10 times and used for calculating response reliability and evaluating model performance (see below). Following data acquisition, recordings were processed using Suite2P^41^ to identify neurons and extract their responses (deconvolved spiking responses). For ANN modelling, data was denoised via singular value decomposition (SVD) to extract stimulus-dependent signals. Specifically, SVD was performed on the ‘trial x time’ matrix for each neuron and the data was projected onto the first singular vector, as we found this closely corresponded to the stimulus-evoked response kernel for the majority of neurons. This denoising was only performed for ANN modelling; validation experiments were analyzed using raw deconvolved spiking responses. To extract stimulus-evoked responses, we averaged the activity in a 700 ms window following stimulus onset and normalized the response for each neuron. In total, we recorded from > 17,500 neurons from 21 animals (V1 = 2,213 cells; LM = 2,315 cells; LI = 3,341 cells; POR = 2,524 cells; AL = 3,214 cells; RL = 4,232 cells), and after taking into account response reliability and explainable variance explained into account (see below), we used > 7,250 model units for our subsequent analyses.

### Response reliability

The reliability of each neuron in response to the presentation of natural images was calculated as the Spearman-Brown corrected correlation coefficient for random half-splits:

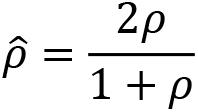

where ρ is the Pearson correlation coefficient, averaged across 100 random samples. For many areas, we found a bimodal distribution of response reliability across all experiments (**Supplementary Figure 1**) and therefore only included neurons with response reliability > 0.5 for subsequent modeling. Reliability could be affected by various factors, including ‘innate’ trial-by-trial variance, non-visual induced activity (e.g. motor movements), or representational drift occurring over the course of a relatively long recording session. Note that RL appeared to be less reliably overall than the other areas, similar to what has been found previously^20^.

### Calculation of explainable variance

For each neuron, the amount of explainable variance was estimated for the 100 repeated stimuli as

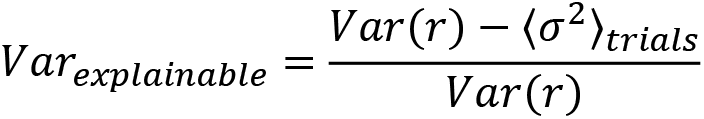

where Var(r) is the variance of the response of a given neuron to all stimuli and 〈σ^2^〉_*trails*_ is the average variance of the responses across repeated trials.

### Linear model

The performance of the ANN model was compared to a simple linear model. Neuron responses were fitted using Partial Least Squares regression with 5-fold cross-validation. Each model was run with a range of bottleneck dimensions (10-20) and the best performing model was chosen.

### Artificial neural network modelling

Deep convolutional neural networks were trained to predict neural responses to natural images. The networks consisted of four blocks, each block composed of 2D convolutional (kernel size = 3, stride = 1), batch normalization, rectified linear unit (ReLU), and max pooling (kernel size = 2, stride = 2) layers, followed by a factorized readout layer^26^. The factorized readout decomposes into independent spatial and feature layers, which consist of tensors of shape (1 x height x width) and (channel x 1 x 1), respectively. The number of channels in the feature layer was set to 512; the height and width of the spatial layer amounted to 8. The spatial layer was further constrained to have non-negative entries and unit Frobenius norm to facilitate its interpretation as a spatial mask. Additionally, L1 regularization was applied to both layers to encourage sparsity.

Networks were trained to predict neural responses to natural images by maximizing the normalized dot product:

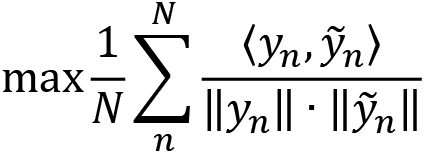

where *y_n_*, *ỹ*_n_ are the response and predicted response of neuron n, respectively, averaged over all neurons N. The networks were trained using Adam optimizer for 30 epochs, with L2 regularization, and early stopping. A hyperparameter search was performed with Oríon (https://orion.readthedocs.io/en/stable/) to find optimal training parameters (**Supplementary Table 1**).

The performance of the model was assessed by calculating the squared Pearson correlation coefficient with a set of held-out data (100 repeated stimuli). A linear regression without offset between the explainable variance and model performance was then performed to estimate the fraction explainable variance explained^23^.

### Spatial mask analysis and alignment

We upscaled the spatial layer of the factorized readout (8 x 8) to the stimulus size (135 x 135). To calculate the area of the spatial mask, we fit a 2D Gaussian

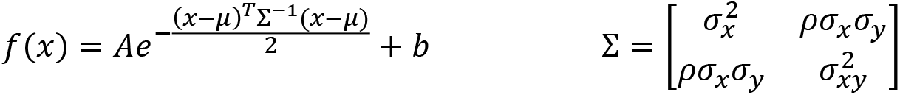

where x are the pixel coordinates, µ the center, σx and σy the standard deviations, ρ the correlation coefficient, A the amplitude, and b the offset. The area of the spatial mask was defined as σxσy of the ellipse at one standard deviation. Only spatial masks that were fit well by the 2D Gaussian (r^2^ > 0.8) were used to estimate the area. For analyses of image statistics, we masked preferred stimuli to include only those regions within two standard deviations of the median spatial mask size of each model. The edges of the mask were further smoothed by applying a Gaussian filter (σ = 5 pixels). We set the background of the preferred stimuli to a pixel value of 0.5.

For *in silico* analyses, the centers of the spatial masks were aligned to the center of the input stimuli. The spatial masks were shifted towards the center with boundaries being wrapped around and values interpolated with 3^rd^ order splines. This was repeated 10 times for more consistent alignment.

### Generation of preferred stimuli

Building on early work that sought to understand the features represented by model units in ANNs^42^, here, preferred stimuli were generated using the Lucent library (https://github.com/greentfrapp/lucent). In brief, starting from random white noise, images were updated to maximize the response of ANN model units by backpropagating the error through the ANN to the input images. To avoid high frequency noise, which is known to result in image artefacts that interfere with interpretability, small random transformations were applied to the images, including padding (0-4 pixels), jitter (0-8 pixels), and rotations (± 10 degrees). The optimization was run for 512 epochs with a learning rate of 3e-3.

### In vivo validation experiments

*In vivo* validation experiments followed the same protocol as described in “2-photon calcium imaging”. After the first recording session, data was analyzed and preferred stimuli were generated for all neurons. For the second recording session, the same field of view was found by aligning the blood vessel patterns. On average, we were able to match ∼80% of neurons across days. Preferred stimuli and up to 500 natural images (randomly selected from ImageNet) were presented three times each. For the quantification, the responses to the three repeats were averaged and normalized to the distribution of responses to natural images for each neuron.

### Distance and partial distance correlation

Distance correlation was used to compare the functional similarity between areas. First, the spatial mask of model units in the ANN models were aligned to the center of the input. Next, 10,000 natural images were presented to the models and the responses were collected and normalized for each neuron. We subsampled the resulting response matrix by randomly choosing 200 neurons and 1,000 stimuli and computed the distance correlation^43^ between all pairs of models, repeated 100 times. The distance correlation was then averaged across repeats. To isolate unique correlations between areas, we computed the partial distance correlation^44^, which approximates the conditional distance correlation^45^:

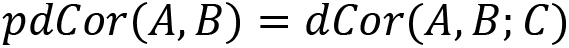

where C is the concatenated matrix over all areas C ≠ A, B.

To visualize the resulting network structure, we converted the pairwise correlation matrix to a dissimilarity matrix d = 1-dCor or d = 1-pdCor and performed multidimensional scaling (MDS) using the scikit-learn library.

### Embeddings of the collection of preferred stimuli

Three image embeddings were used based on (1) perceptual distance, (2) pixel distance, and (3) spatial frequency distance. The distances were defined as the Euclidean distance in the respective embedding spaces. For perceptual distance, we trained a SimCLR model which learns an embedding that is invariant to a custom set of transformations^32,33^. Specifically, we used the backbone of the ResNet18 network and appended a 256-dimensional linear readout layer. The network was then trained to minimize the following loss function:

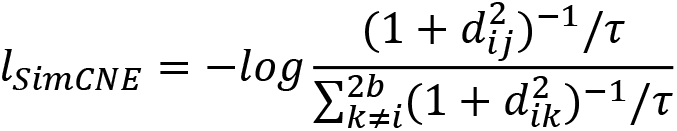

where b=3955 is the batch size and τ = 0.1 a temperature parameter. Setting τ < 1 prevented dimensionality collapse in which the rank of the embedding space would be much smaller than 256 dimensions. d denotes the Euclidean distance calculated for pairs i, j of two random transformations applied to the same preferred image. For invariance to affine transformations, the random transformations were cropping and scaling (area before resizing: 0.08-1 times the total image size of 64 x 64 pixels, resized to 32 x 32 pixels), random rotation (± 90 degree range), random translation (± 0.1 times the total image size), random horizontal and vertical flips, and a Gaussian blur. SimCLR was trained for 500 epochs with Adam optimizer (learning rate = 1e-4) and a cosine annealing schedule for the learning rate. To visualize the SimCLR embedding, UMAP^34^ was used to project the embedding into two-dimensional space (metric = Euclidean, min_dist = 0.1, n_neighbours = 30).

For pixel similarity, the dimensionality of the images was reduced using PCA to match the 256 dimensions of the SimCLR embedding. For the spatial frequency distance, we performed fast Fourier Transform (FFT), calculated the power spectra, removed the DC component, and reduced the dimensionality to 256 using PCA.

A k-nearest neighbours classifier (k=20) was used to assess classification accuracy of the image embeddings (note that the results were qualitatively indistinguishable for k = 10, 20, or 50; data not shown). The classifier was trained on 75% of the data and tested on the remaining 25%. This was repeated 100 times. We separately sampled each area’s preferred stimuli to account for unequal numbers of preferred stimuli generated for each area.

To estimate the overlap between embeddings of preferred stimuli from two areas, we calculated the average distribution of area labels of the 20 nearest neighbours of each image. For example, to calculate the overlap of V1 preferred stimuli with the HVAs, for the 20 nearest neighbours of each preferred stimulus in V1, we counted the number of neighbours that belonged to each HVA. This procedure resulted in a 6 x 6 overlap matrix M for each pair of visual areas. This matrix was not symmetric, so we next averaged across the diagonal, M_symmetric_ = (M+M^T^)/2, i.e. we averaged the overlap of area A with B and the overlap of area B with A. We used this symmetric matrix as a measure for local overlap.

The topological similarity between the local overlap with the functional similarity was computed as the Spearman correlation between the symmetric overlap matrix (**Figure 3d**) and the pairwise partial distance correlation matrix (**Figure 2c**).

### Generating the ‘Feature landscape of mouse visual cortex’

We visualized the two-dimensional UMAP projection of the SimCLR embedding as an image atlas. First, the embedding was scaled to lie within the unit interval [0,1]. Next, we tiled the embedding into an NxN grid (for **Figure 4**, a 40x40 grid is shown). Then, within each tile, we randomly chose an image to display. For visualization purposes, the small number of empty tiles in the 40x40 grid that were fully enclosed within the UMAP projection (i.e. ‘holes’ in the grid) were filled in using the closest image from a neighbouring tile.

### Computing the most representative preferred stimuli

For each preferred stimulus, we defined the degree of representative-ness as the fraction of nearest neighbours (k = 100) in SimCLR’s embedding space that were from the same visual area. We then chose the top 200 most representative stimuli for each region for subsequent analysis in **Figure 5**.

### Preferred stimulus analyses

We computed various low-level image statistics for the preferred stimuli. Pixel intensities ranged from 0 to 1.

1. Luminance was calculated as the average pixel intensity within the spatial mask.
2. To identify dark and light segments, we thresholded stimuli around the background intensity (for dark segments: pixels < 0.49; for light segments: pixels > 0.51). Next, disconnected segments with area > 2 pixels were identified. We analyzed the number of segments and the area, which was normalized by the full area of the stimulus.
3. To calculate the spatial frequency content, we first performed a FFT and averaged the power spectrum radially with bin size = 1 pixel. To calculate the radial average, we converted pixel Cartesian coordinates to polar coordinates, defining the center of the image as the origin (0,0). Pixels were then binned according to their radius, rounded down. Spatial frequency power was normalized for each frequency across all images from all areas.
4. To calculate folio (1-fold) and quarto (2-fold) symmetries, we first performed FFT and averaged the power spectrum axially with bin size = 1/16 π radians. To calculate the axial average, we converted pixel Cartesian coordinates to polar coordinates, defining the center of the image as the origin (0,0). Pixels were then binned according to their angle, rounded down. Next, we defined an n-fold symmetry index (SI) as

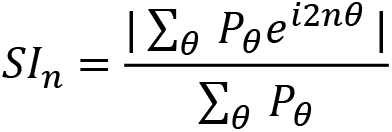

where P_θ_ is the average power at angle θ and the scalar multiplier 2 is due to the inherent point symmetry of the FFT power spectrum. SI is defined on the unit interval with 0 denoting a lack of symmetry and 1 being a fully n-fold symmetric image.

### Data analysis

Statistical significance was assessed using Mann-Whitney U test for unpaired data, Wilcoxon rank sum test for paired data, and Kruskal-Wallis followed by *post hoc* Dunn’s test with Bonferroni correction for multiple comparisons. Data values are reported as mean ± SEM, unless mentioned otherwise. Box plot elements are defined as follows: center line = median, box limits = upper and lower quartiles, whiskers = 1.5 times interquartile range.

### Nomenclature of mouse HVAs

The list^19^ of widely agreed upon mouse HVAs include: LM (lateromedial), AL (anterolateral), RL (rostrolateral), A (anterior), AM (anteromedial), PM (posterior medial), LI (laterointermediate), P (posterior), and POR (postrhinal).

